# Parent–Child Cortical Similarity Indexes Multiple Dimensions of Sensitivity in the Association Between Family Environment and Adolescent Internalizing Symptoms

**DOI:** 10.64898/2026.09.18.752767

**Authors:** Qingyi Li, Ya-Yun Chen, Jungmeen Kim-Spoon, Brooks Casas, Yang Qu, Tae-Ho Lee

**Affiliations:** Department of Psychology, Virginia Tech, Blacksburg, VA, USA; School of Neuroscience, Virginia Tech, Blacksburg, VA, USA; School of Education and Social Policy, Northwestern University, Evanston, IL, USA

**Keywords:** Parent-Child Dyads, Neural Similarity, Environmental Sensitivity, Family Environment, Adolescent Internalizing Symptoms, Brain Morphometry

## Abstract

Family environment is a robust predictor of adolescent internalizing symptoms, yet its strength varies across adolescents. We tested whether this variation is partly captured by how similar an adolescent’s cortical structure is to their parent’s, and whether different forms of this similarity matter in different cortical systems. In 115 parent–child dyads from two cohorts, structural MRI quantified similarity across 400 cortical parcels in two forms: phenotypic similarity in regional morphometry and architectural similarity in interregional morphometric organization. Parent–child similarity was not directly associated with symptoms, whereas its interactions with family environment were jointly significant (χ²(4) = 14.92, *P* = 0.005). Phenotypic similarity in the sensory–limbic system and architectural similarity in the association system each moderated the association between family environment and symptoms (both β = 0.22). In both cases, the association was strongest among less similar dyads. The phenotypic interaction was distributed across sensory–limbic networks and was not reproduced by similarity to unrelated adults or by the child’s cortical atypicality. The architectural interaction was concentrated in the dorsal attention and default mode networks and was distinguishable from unrelated-adult similarity but not from child atypicality. Together, these findings raise the possibility that markers of environmental sensitivity need not reside in the child alone. Parent–child cortical similarity may index a dyadic component of this sensitivity — one not adequately represented by a single similarity score, but distributed across forms of similarity and cortical systems.

**Significance Statement:** Why family environment is more strongly associated with mental health problems in some adolescents than in others is usually explained by characteristics of the child, including temperament, genetic variation, and brain features. Our findings suggest that this sensitivity may also be indexed at the level of the parent–child dyad. Across 115 dyads from two cohorts, family environment was most strongly associated with adolescents’ internalizing symptoms when parent and child had less similar cortical structure. Measures defined on the child alone did not reproduce the full pattern, and the form of similarity that indexed sensitivity differed across cortical systems. Environmental sensitivity may therefore be relational as well as individual, with a dyadic component that is multidimensional rather than singular.

## Introduction

Family environment is among the most robust predictors of adolescent internalizing symptoms across development (1–4). Yet family environment is more strongly associated with internalizing symptoms in some adolescents than in others (5–7). This heterogeneity is commonly conceptualized as environmental sensitivity: the extent to which individuals differ in their responsiveness to environmental experiences. Most studies have examined whether child-level characteristics, including temperament, genetic variation, physiological reactivity, and brain features, moderate associations between environmental experiences and developmental outcomes (7–9). However, recent evidence suggests that this sensitivity may not be a single property. Responsiveness to positive and negative experiences appears to have partly distinct bases (10), suggesting that a single marker may not capture all forms of responsiveness. At the same time, family environment is inherently relational: it unfolds within an ongoing parent–child relationship, whereas candidate markers of sensitivity have typically been measured in the child alone. Sensitivity to family environment may therefore depend not only on characteristics of the child but also on characteristics of the parent–child dyad.

Parent–child neural similarity offers a way to examine these possibilities. Because it is calculated between two brains, it provides a candidate marker at the level of the dyad rather than the child alone. Previous studies have implicated parent–child neural similarity in the relation between family environment and youth adjustment (11–13; for reviews, see 14–17). Other work suggests that different forms of neural resemblance capture partly distinct aspects of intergenerational correspondence (18, 19). Together, these findings raise the possibility that parent–child neural similarity may index multiple components of dyadic sensitivity. Distinguishing such components requires forms of similarity that can be quantified on the same brains, so that differences between them are not confounded with differences in imaging modality.

Cortical morphometry meets this requirement. The same morphometric measurements characterize two aspects of the developing cortex: the features of individual regions and the organization of those features across regions. We refer to parent–child correspondence in these two aspects as phenotypic similarity and architectural similarity, respectively. Phenotypic similarity captures correspondence in regional morphometric features, including cortical thickness, surface area, gray matter volume, and curvature. Each of these features is under common-variant genetic influence, and their genetic architectures differ across the cortex (20, 21). Architectural similarity captures correspondence in the pattern of relations among those regional features. These relations are represented as morphometric similarity networks, in which each connection quantifies the resemblance between two regions’ feature profiles. Morphometric similarity networks have been shown to recapitulate aspects of cortical cytoarchitecture and to correlate with axonal connectivity (22–25). A parent and child may therefore resemble one another in their regional morphometric phenotypes but differ in how those phenotypes are organized across the cortex, or vice versa.

Beyond similarity form, the relevance of each form of similarity may also depend on where in the cortex it is measured. Cortical development follows a broad sensorimotor-to-association axis: compared with earlier-developing systems, association regions mature later, vary more across individuals, and are less constrained by underlying anatomy (26–28). Regional morphometric features and interregional organization also follow partly distinct developmental courses. Regional features continue to change across adolescence, particularly in association cortex (26, 27, 29), whereas interregional organization reflects coordinated maturation among regions and is reorganized across adolescence in spatially varying directions (23, 30). These developmental differences motivate testing whether the association between family environment and symptoms is moderated by the same form of similarity across cortical systems or by different combinations of form and system.

Across two cohorts of parent–child dyads, we used structural MRI to quantify phenotypic similarity from regional morphometric profiles (31) and architectural similarity from morphometric similarity networks (24). Both forms were quantified for the whole cortex and within two systems spanning this developmental axis: a sensory–limbic system comprising the visual, somatomotor, and limbic networks, and an association system comprising the dorsal attention, salience/ventral attention, frontoparietal, and default mode networks (Fig. 1). The limbic network was grouped with the sensory rather than the association networks, and the two systems thus contrast heteromodal association cortex with the rest of the cortex. Because paralimbic cortex follows a maturational trajectory distinct from neocortex as a whole (30), the limbic network was also analyzed on its own (SI Appendix, Table S4b). We tested (a) whether parent–child cortical similarity moderated the association between family environment and internalizing symptoms, with variation in the strength of this association serving as the operational index of sensitivity; (b) whether the moderation varied across combinations of similarity form and cortical system; and (c) whether any observed moderation was specific to similarity with the child’s own parent or could be reproduced by similarity to unrelated adults or by the child’s own cortical atypicality.

**Fig. 1.**
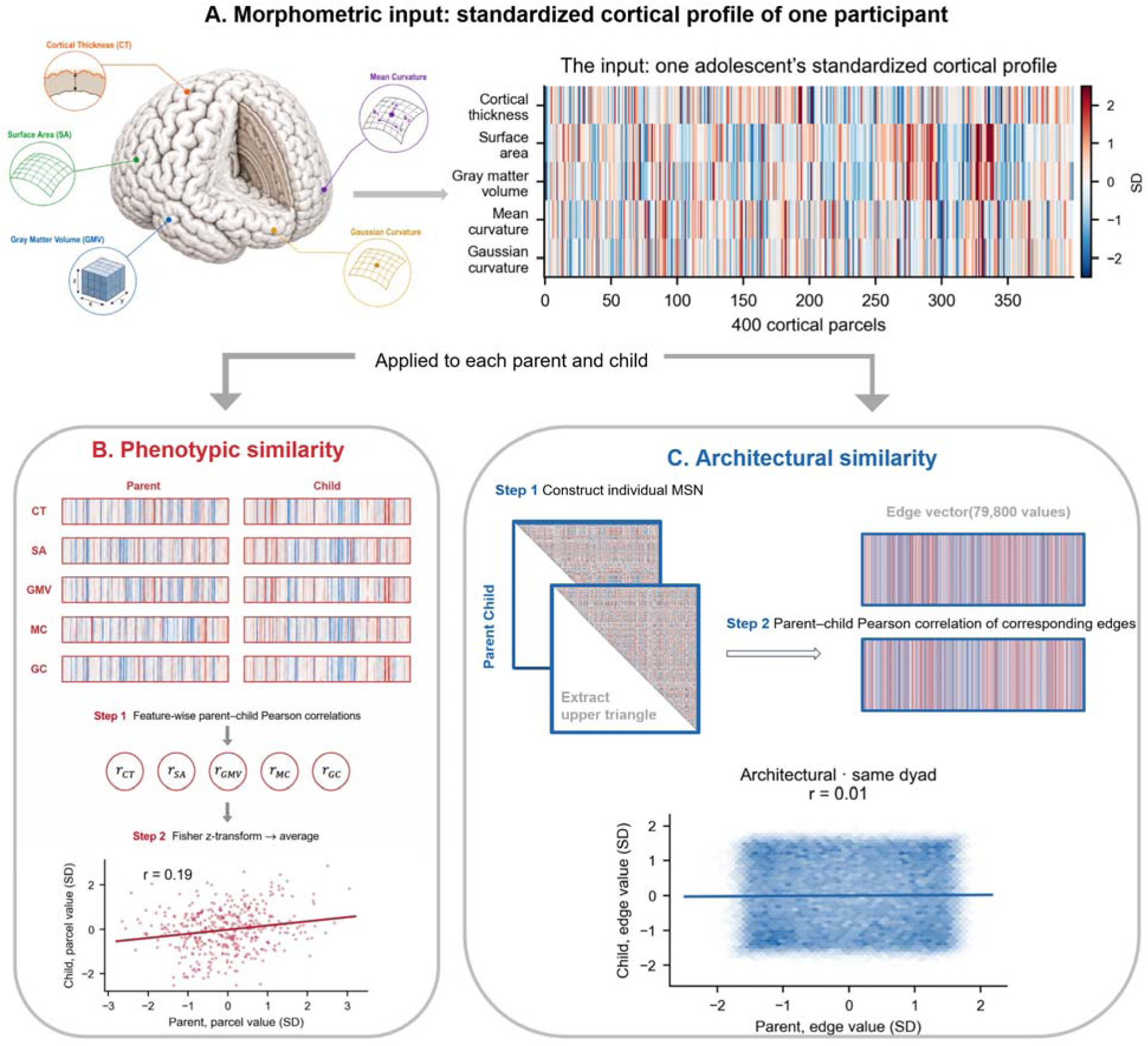
Two forms of parent–child structural brain similarity, computed from the same data. (A) Five morphometric features per Schaefer-400 parcel — cortical thickness, surface area, gray-matter volume, mean curvature, and Gaussian curvature — standardized as deviations from the group-typical pattern (*SI Appendix*, SM5); the heatmap shows one adolescent. (B) Phenotypic similarity: parent and child profiles correlated across parcels per feature and averaged on the Fisher-z scale (example dyad, cortical thickness, *r* = 0.19). (C) Architectural similarity: each participant’s morphometric similarity network (24), with the parent’s and child’s standardized edge vectors (79,800 edges) correlated (same dyad, *r* = 0.01). The two formss can dissociate within a dyad and are entered as separate moderators. Red denotes the phenotypic and blue the architectural similarity throughout.

## Results

### Parent–child similarity was higher than permuted dyads

We first tested whether the similarity scores distinguished true parent–child dyads from cross-family pairings. For each network and similarity form, similarity to the child’s own parent was compared with a null distribution generated by pairing the same children with parents from other families across 10,000 permutations (Fig. 2B; *SI Appendix*, Table S5c). These comparisons used all 120 dyads with usable imaging. All comparisons were significant except architectural similarity in the limbic network, which did not differ from the null distribution in either Cohort 1 (*P* = 0.29) or Cohort 2 (*P* = 0.17). At the whole-cortex level the same comparison is shown as distributions (Fig. 2C). We next tested whether each child’s own parent could be identified among parents from other families. For each child the own parent was ranked against every other family’s parent (Fig. 2D). At the whole-cortex level the own parent ranked first in 51.0% (phenotypic) and 37.3% (architectural) of comparisons in Cohort 1 and in 52.2% and 31.9% in Cohort 2, against chance rates of 2.0% and 1.4% (all *Ps* < 0.0001).

**Fig. 2.**
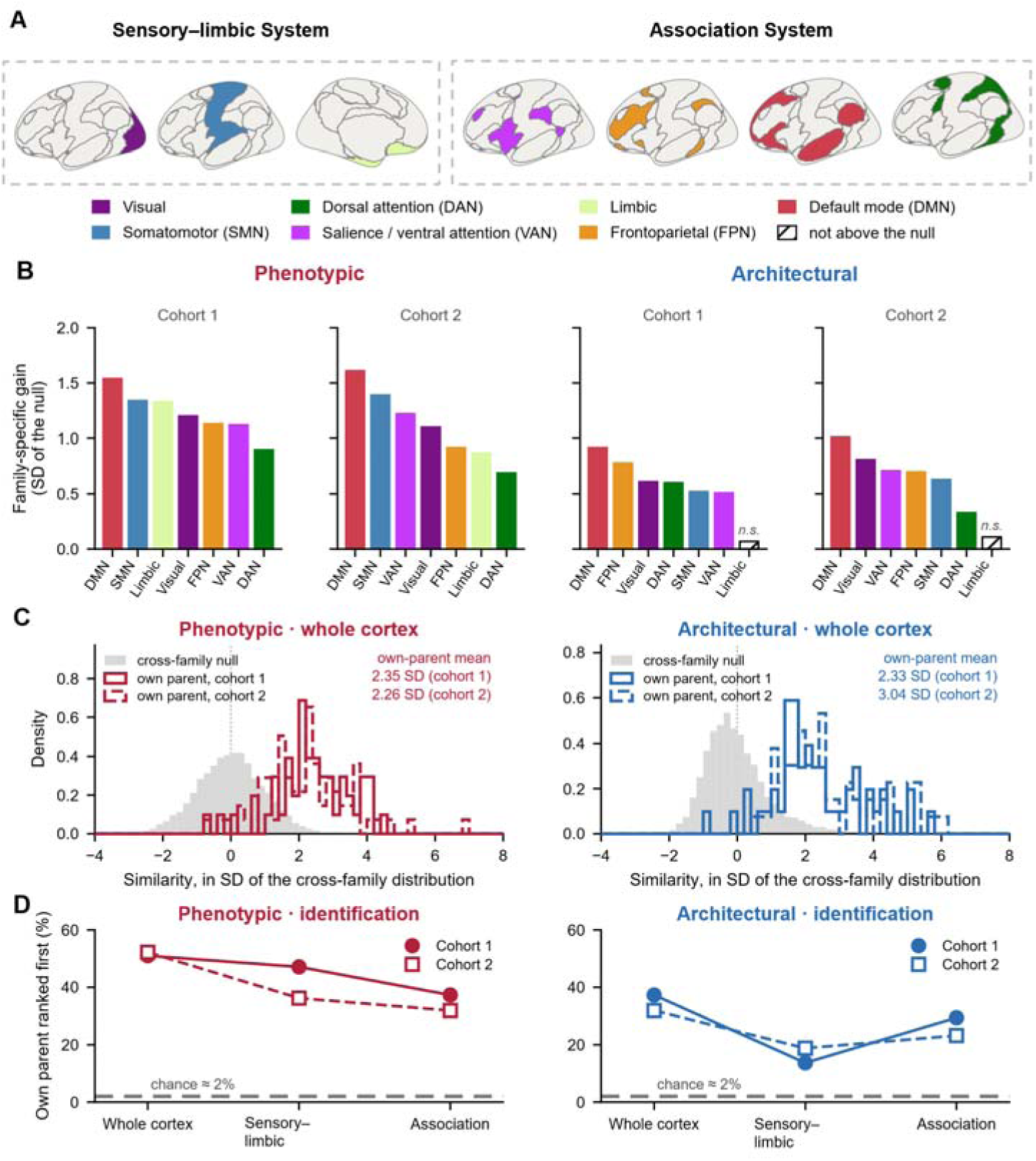
Parent–child similarity carries family-specific information, and where in the cortex it does. (A) The seven networks (49) and the two systems used in the analyses. (B) Family-specific gain per network: how far real dyads exceed the same children paired with every other family’s parent, in SD of that cross-family distribution, ranked within cohort. Filled bars exceed their permutation null at *P* < 0.05 (10,000 draws); hatched bars do not — the limbic network at the architectural similarity in both cohorts (*P* = 0.29 and 0.17; *SI Appendix*, Table S5c). (C) The same quantity for the whole cortex; own-parent means are 2.35 and 2.26 SD phenotypically and 2.33 and 3.04 SD architecturally (Cohorts 1 and 2). (D) Proportion of children whose own parent ranks first against every other family’s parent; chance is 2.0% and 1.4% (dashed rule) and all points exceed it (*SI Appendix*, Table S5d) (*P* < 0.0001). Panels B–D use all 120 dyads with usable imaging (51 in Cohort 1 and 69 in Cohort 2); the moderation models in Figs. 3 and 4 use the 115 dyads with complete questionnaire data.

### Phenotypic and architectural similarity moderated the environment–symptom association in distinct cortical systems

Parent–child similarity was not associated with internalizing symptoms. Across the six similarity measures defined by three cortical units (whole cortex, sensory–limbic system, and association system) and two similarity forms (phenotypic and architectural), standardized coefficients ranged from −0.03 to −0.15, and none was statistically significant (all *Ps* ≥ 0.06; *SI Appendix*, Table S1; Fig. 3A, left). Family environment itself was associated with symptoms in both cohorts and in the pooled sample (β = −0.43, *P* < 0.0001; Cohort 1 β = −0.35, *P* = 0.043; Cohort 2 β = −0.48, P < 0.0001; *SI Appendix*, Table S1).

**Fig. 3.**
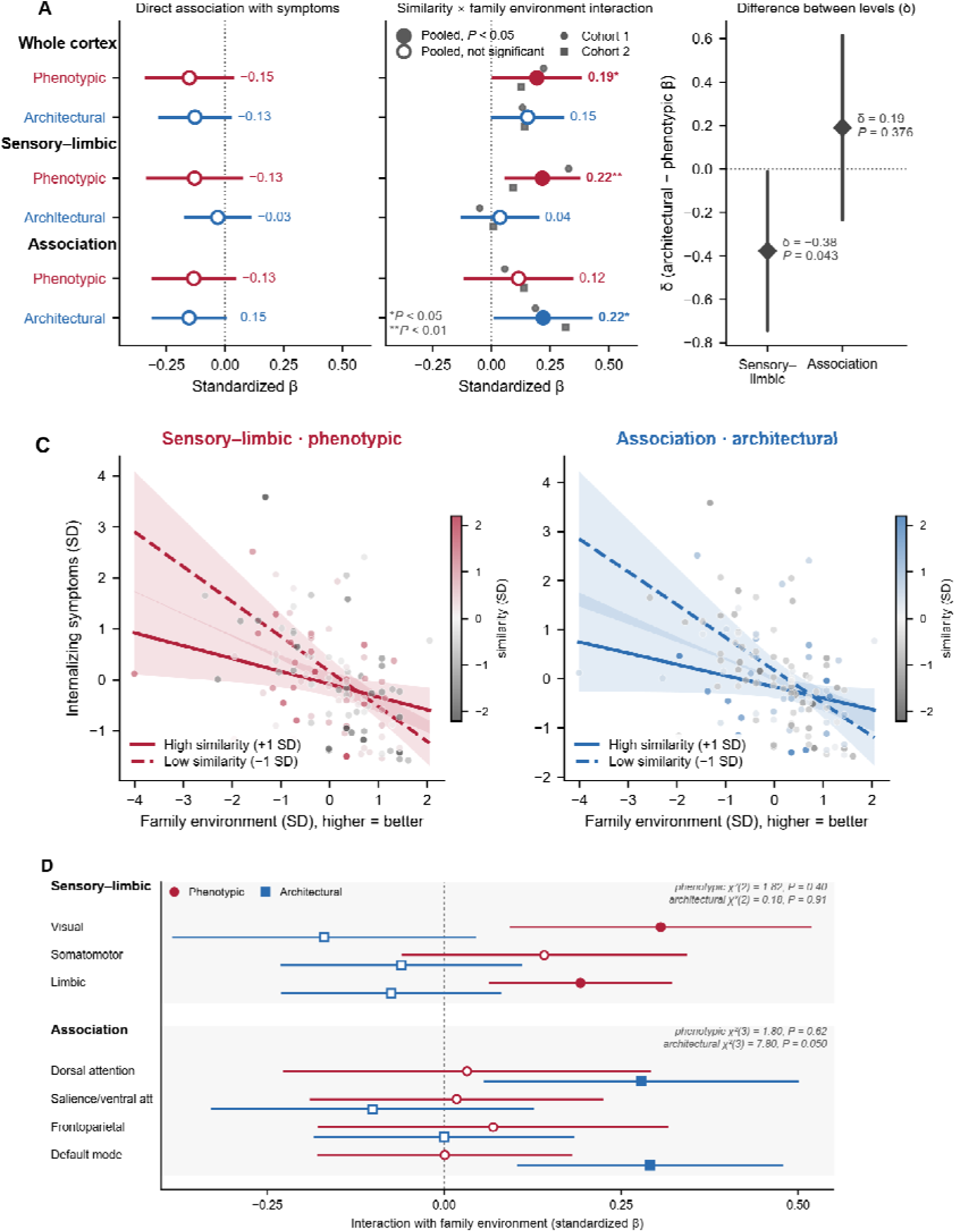
Parent–child similarity indexes sensitivity to family environment. (A) The six system-by-form scores entered two ways: their direct association with symptoms (left; all estimates −0.03 to −0.15, *P* ≥ 0.06) and their interaction with family environment (right). Filled circles, pooled *P* < 0.05; open circles, not significant; bars, 95% CIs; small grey markers, per-cohort estimates. \**P* < 0.05, \*\**P* < 0.01, uncorrected. The four interactions are jointly nonzero (Wald χ²(4) = 14.92, *P* = 0.005). (B) The similarity-form difference within each system, δ = β(architectural × E) − β(phenotypic × E), from a single model containing both scores. (C) The environment–symptom association at ±1 SD of the moderator with 95% bands, for the significant cell at each form; points are all dyads, shaded by similarity. (D) The same interaction at each of the seven networks within its system, ordered along the sensorimotor-to-association axis; filled markers, *P* < 0.05 uncorrected.

Across the whole cortex, the interactions at both similarity forms were of borderline significance (phenotypic β = 0.19, *P* = 0.049; architectural β = 0.15, *P* = 0.054; *SI Appendix*, Table S2b) and are treated as secondary. We then examined the two pre-specified systems: the sensory–limbic system and the association system (Fig. 2A).

The two forms of similarity showed moderation in different systems (Fig. 3A, *SI Appendix*, Table S1). Phenotypic similarity in the sensory–limbic system moderated the association between family environment and symptoms (β = 0.22, *P* = 0.009), whereas architectural similarity in the same system did not (β = 0.04, *P* = 0.670). In the association system the pattern was reversed: architectural similarity moderated the association (β = 0.22, *P* = 0.041) and phenotypic similarity did not (β = 0.12, *P* = 0.330). Because the two forms are computed on the same dyads from the same parcels, their difference was estimated from a single model containing both (Fig. 3B). Within the sensory–limbic system the two forms differed (δ = −0.38, *SE* = 0.19, *P* = 0.043); within the association system they did not (δ = 0.19, *SE* = 0.21, *P* = 0.376). With all four system-by-form scores entered together, the four interactions were jointly different from zero (Wald χ²(4) = 14.92, *P* = 0.005), with no variance inflation factor among the interaction terms exceeding 1.91 and dyad-level correlations among the four scores ranging from 0.16 to 0.55 (*SI Appendix*, Table S2c), indicating that the four scores carry distinct rather than redundant information. In that model the difference between similarity forms reversed in sign across the two systems (δ = −0.42 in the sensory– limbic system and +0.30 in the association system), and the difference between these two differences was itself significant (Δ = −0.72, SE = 0.29, *t*(109) = −2.50, *P* = 0.014; *SI Appendix*, Table S1).

Because higher family-environment scores indicated better relationships whereas higher outcome scores indicated more symptoms, the environment–symptom association was negative. Both significant interactions reflected stronger associations between family environment and internalizing symptoms among less similar dyads, with the environment–symptom association attenuating as parent–child similarity increased. For sensory–limbic phenotypic similarity, the simple slope of family environment was β = −0.68 (*P* < 0.001) at one standard deviation below the mean of similarity and β = −0.25 (*P* = 0.011) at one standard deviation above it. For association architectural similarity, the corresponding slopes were β = −0.67 (*P* < 0.001) and β = −0.23 (*P* = 0.049), respectively (Fig. 3C).

This result was stable under alternative network groupings. Under two divisions defined in advance, the difference between similarity forms was present in the visual and somatomotor networks alone (δ = −0.43, *P* = 0.014) and in the limbic network alone (δ = −0.27, *P* = 0.012), but never in the association system (all *P*s ≥ 0.37); dyadic specificity was in the same direction under these divisions, although not every contrast reached significance (*SI Appendix*, Table S4b). The two cohorts were pooled rather than treated as replications: five of the six similarity scores showed interactions in the same direction in both cohorts, but neither cohort was powered to detect them alone, so the per-cohort estimates are directionally consistent with the pooled results rather than independent replications (*SI Appendix*, Table S2b).

### Dyadic specificity of moderation

To test whether the moderation was specific to similarity with the child’s own parent, we evaluated two non-dyad-specific comparison measures using the same Brain × Environment model. Unrelated-adult similarity was defined as the child’s mean similarity to parents from other families and indexed general resemblance to adult cortex (Fig. 4A). Child atypicality quantified the magnitude of the child’s deviation from the group-average cortical profile (Fig. 4A). Together, these controls tested whether the apparent dyadic moderation could instead be attributed to characteristics of the child — general resemblance to adult cortex or cortical atypicality — rather than correspondence with the child’s own parent.

**Fig. 4.**
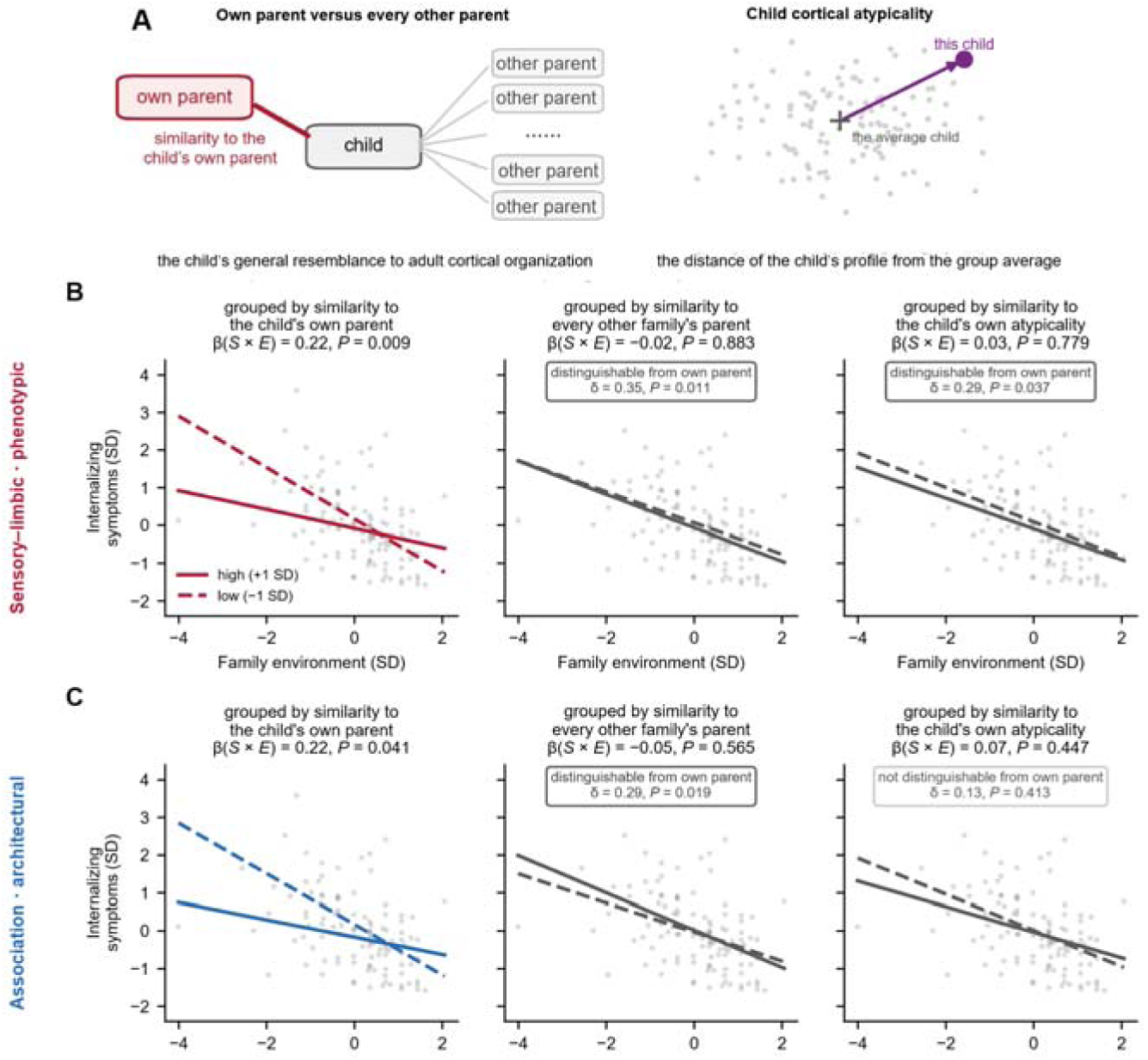
The moderation is specific to the child’s own parent, established at the phenotypic similarity. (A) The two control quantities: the child’s mean similarity to every other family’s parent (general resemblance to adult cortical organization) and child cortical atypicality (the distance of the child’s standardized profile from the group average). (B, C) The environment–symptom association fitted at ±1 SD of each candidate moderator, for sensory–limbic phenotypic similarity (B) and association architectural similarity (C); the left column reproduces the own-parent moderation from Fig. 3C, and grey points are identical across panels — only the grouping variable changes. Panel headers give each score’s own interaction; boxes give the contrast δ = β(own parent × E) − β(control × E) from a joint model. n = 115 dyads; covariates and clustering as in Methods. Full contrasts: *SI Appendix*, Table S3a.

Dyadic specificity requires three things: similarity to the child’s own parent moderates the environment–symptom association, the two child-level alternatives do not, and the differences are themselves significant. Each difference (δ) was therefore tested directly, with both scores in one model (32).

For sensory–limbic phenotypic similarity, all three conditions held (Fig. 4B). Similarity to the child’s own parent moderated the association (β = 0.22, *P* = 0.009); similarity to unrelated parents did not (β = −0.02, *P* = 0.883), nor did child atypicality (β = 0.03, *P* = 0.779); and both differences were significant (δ = 0.35, *P* = 0.011 against the unrelated adults; δ = 0.29, *P* = 0.037 against child atypicality). Across dyads the own-parent scores were nearly uncorrelated with both control measures in every unit-by-form cell (|*r*| ≤ 0.14; *SI Appendix*, Table S3a), so each contrast compares quantities that share almost no variance, and the own-parent estimate changed little when either control was entered into the model.

In the association system (Fig. 4C), architectural similarity to the child’s own parent moderated the association (β = 0.22, *P* = 0.041); the difference from unrelated-parent similarity was significant (δ = 0.29, *P* = 0.019), whereas the difference from child atypicality was not (δ = 0.14, *P* = 0.413). As in the sensory–limbic cell, the own-parent estimate changed little when either control was entered (β = 0.221 and 0.223, against 0.220 alone), and child atypicality itself showed a weaker interaction that did not differ reliably from it. The pre-specified specificity family comprised the three scores carrying an interaction — whole-cortex phenotypic, sensory–limbic phenotypic, and association-system architectural — each contrasted with both controls, six tests in all. Five survived Benjamini–Hochberg correction (adjusted *P* ≤ 0.044), including both whole-cortex phenotypic contrasts (δ = 0.37 and 0.30); the contrast between association-system architectural similarity and child atypicality was not (*SI Appendix*, Table S3a).

Exploratory network-level analyses suggested that phenotypic moderation was broadly distributed across the sensory–limbic networks, whereas architectural moderation was concentrated in the dorsal attention and default mode networks. At the association-system level, the own-parent architectural interaction differed from the unrelated-adult interaction but not from the child-atypicality interaction. Full network-specific interaction estimates and specificity contrasts are reported in the *SI Appendix*, Tables S5a and S3b.

### Robustness and validation analyses

All estimates were re-derived under two alternative covariate specifications, under which both primary interactions and the contrasts establishing dyadic specificity were similar in magnitude. A secondary strategy that combined cohort-specific estimates by inverse-variance weighting rather than pooling dyads shifted their relative strength toward the association-system architectural effect. These are alternative analyses of the same data rather than independent tests, and all are reported in full in *SI Appendix*, Tables S2a, S2b, and S3a; whole-cortex and exploratory findings varied more and are treated as secondary.

Network-level heterogeneity in association-system architectural moderation was χ²(3) = 7.80, *P* = 0.050 (Fig. 3D). Interaction estimates were positive in the dorsal attention (β = 0.25) and default mode (β = 0.24) networks and near zero in the salience/ventral attention and frontoparietal networks (both β = −0.01; *SI Appendix*, Table S4a). Other exploratory network-by-measure findings varied across analytic strategies and are not interpreted individually (*SI Appendix*, Table S5b).

As an additional biological validation, regional parent–child correspondence was modestly associated with published estimates of SNP heritability (21) for both similarity forms (phenotypic composite, ρ = 0.25, spin-test *P* = 0.001; architectural, ρ = 0.21, *P* < 0.001). Both associations remained after adjustment for the sensorimotor–association axis (*SI Appendix*, SM9). This alignment is expected under any account with a genetic component and so does not separate genetic from shared-environmental contributions; nor does it bear on whether the moderation exceeds child-level information, a separate question that the atypicality contrasts above address only in part.

## Discussion

In 115 parent–child dyads, cortical similarity indexed multidimensional variation in how strongly family environment was associated with adolescent internalizing symptoms. Sensitivity to the family environment was therefore partly a property of the dyad rather than of the child alone, and it was not captured by a single similarity score: whether similarity carried the moderation depended on its form and on the cortical system in which it was measured — phenotypic in sensory–limbic cortex, architectural in association cortex.

The findings support a dyadic interpretation in two respects. First, environmental sensitivity has generally been indexed using characteristics of the individual child (33); here, the moderator was defined jointly by child and parent, so child-level measures may miss part of the variation in how strongly family environment relates to internalizing symptoms. Second, the moderation was not a matter of how adult-like the child’s cortex was: similarity to unrelated adults did not reproduce it in either system, and its interaction coefficients differed from those for the child’s own parent (both δ ≥ 0.29, *P* ≤ 0.019). Whether the moderation also exceeds the child’s own cortical atypicality was not stably resolved at this sample size: the contrast favored separability for the sensory– limbic phenotypic effect but hovered near the significance threshold across analytic choices, and did not favor it for the association architectural effect (δ = 0.14, 95% CI [−0.19, 0.46]) — an interval extending from no difference to a difference twice the size of the own-parent interaction itself, so the contrast is uninformative rather than negative at this sample size. The alternative this leaves open is substantive rather than residual: cortical similarity networks are heritable at the level of individual edges, with genetic effects organized along cortex-wide gradients that overlap genetically with neuropsychiatric traits (34), so a child’s architectural atypicality may itself index partly heritable liability. The direction of the moderation is nonetheless difficult to derive from child-level transmission alone: the environment–symptom association was strongest among the least similar dyads, and dissimilarity from one’s own parent is a relational quantity rather than a characteristic of the child. This pattern is consistent with family systems perspectives in which the parent–child relationship, rather than the individual, is the unit of developmental analysis (35). The own-parent correspondence itself may reflect biological and interactional processes accumulating within that particular family — genetic transmission, shared household environments, and their interplay across development — although these sources cannot be separated here.

The present findings also indicate that environmental sensitivity is not adequately represented by a single dyadic similarity score. The same family environment–symptom association was moderated by different forms of parent–child similarity depending on the form and cortical system in which it was measured. Evidence for partially separable components of environmental sensitivity has emerged along other axes: twin analyses identified partly distinct heritable components of responsiveness to negative and positive experiences (10), and evidence for sensitivity varies across temperament markers (36). The present results add two further axes within a single dyadic neural framework: the similarity form and the cortical system at which parent–child correspondence is measured.

These dimensions are grounded in established features of cortical organization. Phenotypic similarity captures correspondence in the morphometric characteristics of cortical regions, whereas architectural similarity captures correspondence in the relations among those regions, an organization linked to cytoarchitectonic divisions and axonal connectivity (24, 37). Cortical systems likewise differ developmentally: sensory and paralimbic territories follow trajectories distinct from association cortex, which develops over a more prolonged period and shows greater interindividual variability (26–28, 30, 38). The observed pattern of dyadic sensitivity thus maps onto established dimensions of cortical organization.

Why sensitivity took this shape— phenotypic in the sensory–limbic system and architectural in the association system —may depend on the joint properties of form and system. The two forms differ in their relative openness to experience: regional features are the more malleable of the two, continuing to change across adolescence, whereas interregional organization is more strongly anchored in cytoarchitecture and connectivity (24, 37). The two systems differ in how far development individualizes them, association cortex most of all (38–40). Moderation may accordingly have appeared in the crossed pairings — the more malleable form within the constrained system, and the more anchored form within the individualized system — while each matched pairing was null for a different reason. Where form and system are both constrained, family-specific variation appears to reach a floor: architectural similarity in the sensory–limbic system showed the weakest own-parent advantage of any combination, with none detectable in the limbic network (Fig. 2B), leaving little dyad-specific variation to index. Where both are open, familial correspondence persists but may lose its purchase: phenotypic correspondence in association cortex remained strongly family-specific — the default mode network showed the largest own-parent advantage of any network (Fig. 2B) — yet carried no moderation, consistent with regional features being continually reworked by the child’s own individualization. On this explanation, parent–child similarity indexes sensitivity where a familial pattern is both preserved and still has room to matter. This account was formulated after the results were known and cannot be tested with cross-sectional data; it predicts, however, that the relative contributions of the two forms should shift across development, a possibility that longitudinal dyadic imaging could test. Architectural moderation was concentrated in the dorsal attention and default mode networks, with near-zero estimates in the frontoparietal and salience/ventral attention networks. Although association networks show substantial and partly heritable individual variation (39, 40), such variation alone cannot explain this localization because other association networks showed little moderation. The dorsal attention and default mode networks are implicated in externally and internally directed cognition, respectively (41–43), but whether these functional roles explain their involvement cannot be determined from structural data. Because this localization was identified post hoc, it requires independent replication.

Both interactions showed the same form of moderation. The association between family environment and symptoms was stronger at lower levels of similarity (β = −0.68 and −0.67 at 1 SD below the mean) and weaker at higher levels of similarity (β = −0.25 and −0.23 at 1 SD above the mean; Fig. 3C). Lower parent–child similarity therefore indexed greater contingency of adolescents’ symptoms on current family conditions, rather than higher symptom levels in itself. One possibility is that lower similarity reflects accumulated divergence in the developmental experiences of parents and children (41), although the present design cannot determine how this correspondence developed or why it moderated the association. The direction is consistent with a prior longitudinal study in which greater parent–adolescent functional similarity attenuated the association between household chaos and youth adjustment (12). We therefore interpret the result as attenuation of the environment–symptom association rather than as a protective effect of similarity.

Taken together, these findings provide preliminary evidence that environmental sensitivity may be partly dyadic and multidimensional. Parent–child cortical similarity indexed sensitivity at the dyadic level, and this association differed across forms of similarity and cortical systems. Five limitations should be noted. (a) The sample of 115 dyads — large for dyadic imaging, but small for estimating interactions — yielded significant primary interactions only in the pooled sample, with the single exception of the association-system architectural interaction in Cohort 2 (*SI Appendix*, Table S2b); the cohorts also differed in instruments, scanners, and sociodemographic composition. The pooled estimate therefore rests on directional consistency across cohorts rather than on independent replication, and effect sizes may overestimate population values (42–44); the pre-specified, aggregated nature of the similarity scores limits, but does not remove, this concern (45). (b) The analyses were not preregistered; alternative specifications and both cross-cohort estimators are reported (*SI Appendix*). (c) The cross-sectional design permits neither causal nor developmental inference, and genetic transmission cannot be separated from shared environment; longitudinal dyadic imaging would be needed to test whether similarity tracks changing family conditions, and genetically informed designs to separate transmission pathways. (d) Leave-one-family-out standardization may yield larger deviation variance in more variable networks (39, 40), which could inflate the apparent prominence of specific networks; within the sensory–limbic system the numerically largest contribution came from the visual network, although power to resolve within-system differences was limited. (e) Both cohorts came from a single national context, leaving cross-cultural generalizability unknown.

## Materials and Methods

### Participants

The primary analytic sample comprised 115 parent–child dyads from two separately recruited U.S. cohorts: 48 dyads from 43 families in Cohort 1 (children aged 8–17 years) and 67 dyads from 67 families in Cohort 2 (children aged 13–17 years). Imaging-only analyses included all dyads with usable MRI data (51 and 69, respectively). Some Cohort 1 families contributed sibling dyads; analyses accounted for this dependence by clustering standard errors by family and excluding same-family parents from cross-family comparisons. Parents provided written informed consent and minors provided assent; the protocol was approved by the Virginia Tech Institutional Review Board. Recruitment, exclusion chains, missing-data handling, and family composition are detailed in Table 1 and *SI Appendix*, SM1.

**Table 1.**
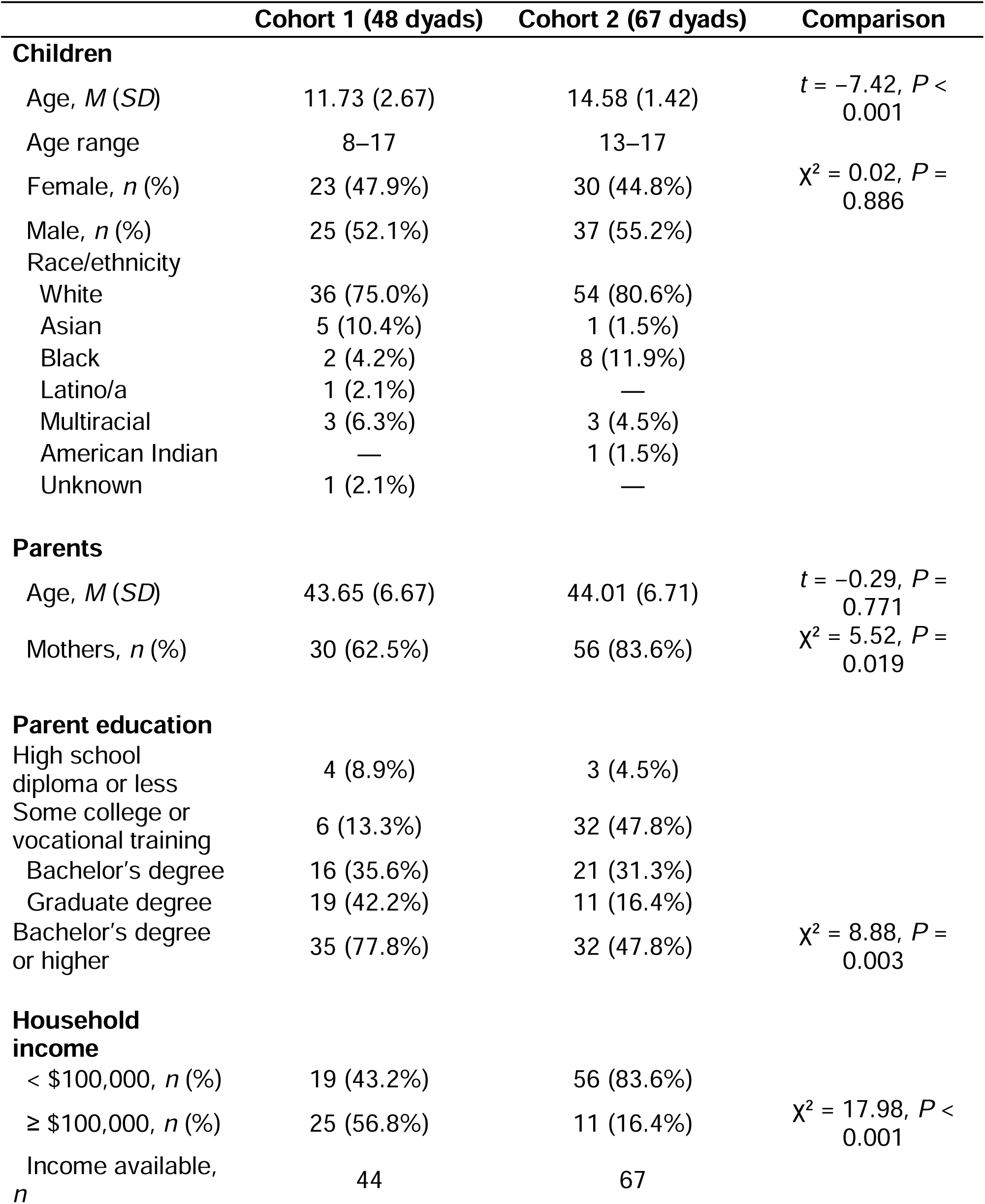
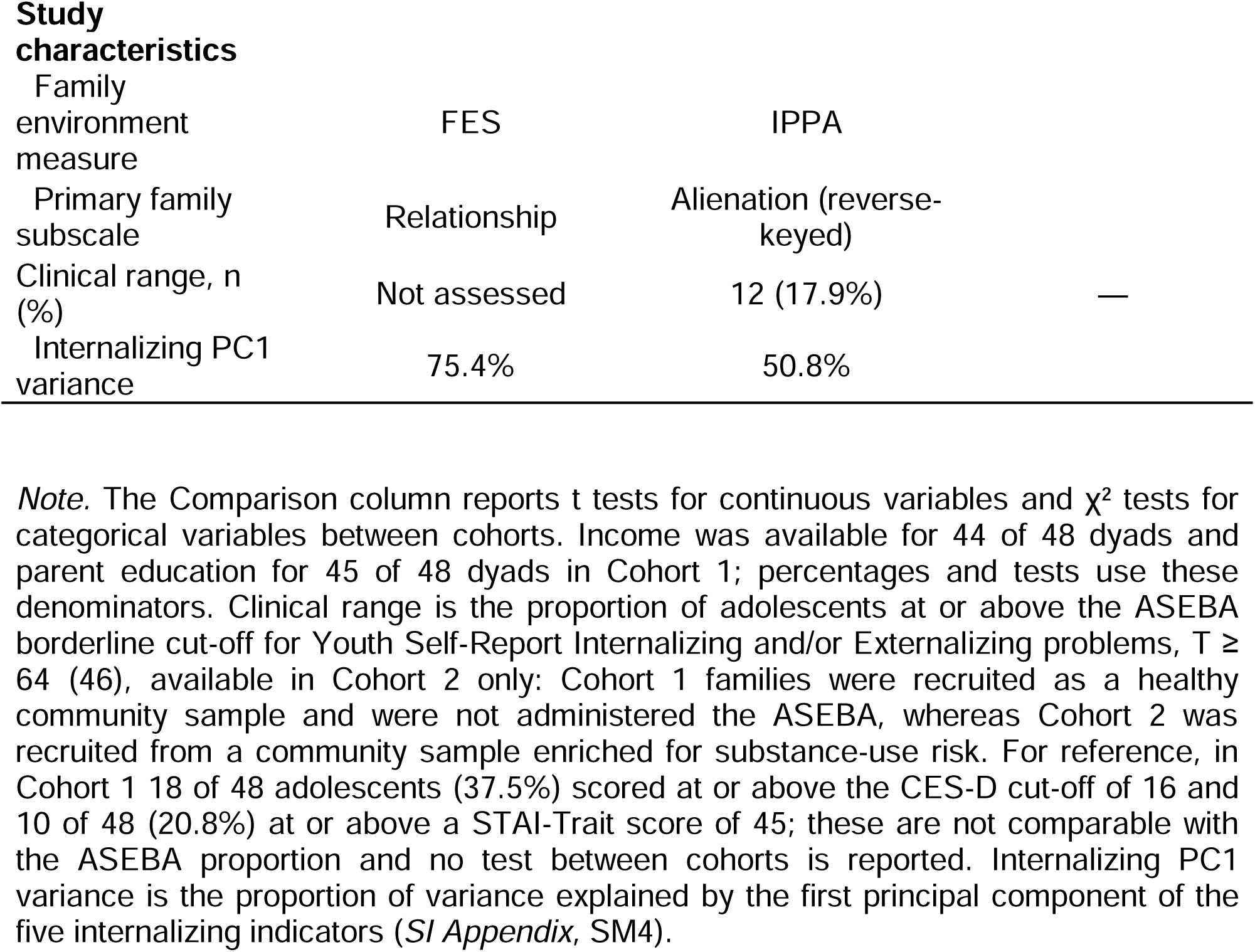
Sample characteristics across two cohorts.

|  | **Cohort 1 (48 dyads)** | **Cohort 2 (67 dyads)** | **Comparison** |
| --- | --- | --- | --- |
| **Children** |  |  |  |
| Age, *M* (*SD*) | 11.73 (2.67) | 14.58 (1.42) | $t = -7.42, P < 0.001$ |
| Age range | 8–17 | 13–17 |  |
| Female, *n* (%) | 23 (47.9%) | 30 (44.8%) | $\chi^2 = 0.02, P = 0.886$ |
| Male, *n* (%) | 25 (52.1%) | 37 (55.2%) |  |
| Race/ethnicity |  |  |  |
| White | 36 (75.0%) | 54 (80.6%) |  |
| Asian | 5 (10.4%) | 1 (1.5%) |  |
| Black | 2 (4.2%) | 8 (11.9%) |  |
| Latino/a | 1 (2.1%) | — |  |
| Multiracial | 3 (6.3%) | 3 (4.5%) |  |
| American Indian | — | 1 (1.5%) |  |
| Unknown | 1 (2.1%) | — |  |
| **Parents** |  |  |  |
| Age, *M* (*SD*) | 43.65 (6.67) | 44.01 (6.71) | $t = -0.29, P = 0.771$ |
| Mothers, *n* (%) | 30 (62.5%) | 56 (83.6%) | $\chi^2 = 5.52, P = 0.019$ |
| **Parent education** |  |  |  |
| High school diploma or less | 4 (8.9%) | 3 (4.5%) |  |
| Some college or vocational training | 6 (13.3%) | 32 (47.8%) |  |
| Bachelor's degree | 16 (35.6%) | 21 (31.3%) |  |
| Graduate degree | 19 (42.2%) | 11 (16.4%) |  |
| Bachelor's degree or higher | 35 (77.8%) | 32 (47.8%) | $\chi^2 = 8.88, P = 0.003$ |
| **Household income** |  |  |  |
| < \$100,000, *n* (%) | 19 (43.2%) | 56 (83.6%) | |
| ≥ \$100,000, *n* (%) | 25 (56.8%) | 11 (16.4%) | $\chi^2 = 17.98, P < 0.001$ |
| Income available, *n* | 44 | 67 |  |
| **Study characteristics** |  |  |  |
| Family environment measure | FES | IPPA |  |
| Primary family subscale | Relationship | Alienation (reverse-keyed) |  |
| Clinical range, n (%) | Not assessed | 12 (17.9%) | — |
| Internalizing PC1 variance | 75.4% | 50.8% |  |

*Note.* The Comparison column reports t tests for continuous variables and $\chi^2$ tests for categorical variables between cohorts. Income was available for 44 of 48 dyads and parent education for 45 of 48 dyads in Cohort 1; percentages and tests use these denominators. Clinical range is the proportion of adolescents at or above the ASEBA borderline cut-off for Youth Self-Report Internalizing and/or Externalizing problems, $T \geq 64$ (46), available in Cohort 2 only: Cohort 1 families were recruited as a healthy community sample and were not administered the ASEBA, whereas Cohort 2 was recruited from a community sample enriched for substance-use risk. For reference, in Cohort 1 18 of 48 adolescents (37.5%) scored at or above the CES-D cut-off of 16 and 10 of 48 (20.8%) at or above a STAI-Trait score of 45; these are not comparable with the ASEBA proportion and no test between cohorts is reported. Internalizing PC1 variance is the proportion of variance explained by the first principal component of the five internalizing indicators (*SI Appendix*, SM4).

#### Imaging and morphometry

T1-weighted images (3T) were processed with FreeSurfer 8.1.0 (47). The cortex was parcellated into 400 parcels (48) organized into seven canonical networks (49), and five morphometric features were extracted per parcel: cortical thickness, surface area, gray-matter volume, Gaussian curvature, and mean curvature (*SI Appendix*, SM2).

### Parent–child similarity

Before any comparison, three standardization steps removed sources of resemblance not specific to a family: head size (leave-one-family-out eTIV residualization), the overall magnitude of each participant’s cortical profile (within-subject z-scoring across parcels), and the population-average anatomical pattern (leave-one-family-out cross-participant standardization). Phenotypic similarity is the parent–child Pearson correlation of parcel-wise values per feature, combined into a Fisher-z composite; architectural similarity is the correlation between parent and child morphometric similarity network edge vectors (24, 50). Scores were computed for the whole cortex, for a sensory–limbic system (visual, somatomotor, limbic) and an association system (dorsal attention, salience/ventral attention, frontoparietal, default mode), and for each network (*SI Appendix*, SM5–SM7).

### Measures

Family environment was the adolescent’s report of family relationship quality: the FES Relationship subscale in Cohort 1 (51) and the reverse-scored IPPA Alienation subscale in Cohort 2 (52). Internalizing symptoms were the first principal component of five standardized indicators per cohort (*SI Appendix*, SM3–SM4).

### Statistical analysis

Brain × Environment models regressed internalizing symptoms on similarity, family environment, and their interaction, adjusting for parent–child age difference, sex pairing, child race, and cohort, with variables z-standardized within cohort and standard errors clustered by family. The primary strategy pooled dyads across cohorts; a secondary strategy combined cohort-specific estimates by inverse-variance weighting. Differences between similarity form and dyadic specificity (against unrelated-parent similarity and child cortical atypicality) were tested as formal contrasts (δ) from joint models. Whether the difference between forms itself differed between the two cortical systems was tested as a further contrast (Δ) in the model containing all four system-by-form scores, with the standard error taken from the full covariance matrix of the four interaction coefficients. Every model was re-estimated under two alternative covariate specifications (*SI Appendix*, SM8) and under the secondary inverse-variance strategy. These are reported in full in the *SI Appendix* and are treated as sensitivity analyses of the same data rather than as independent tests of the primary claims.

### Use of generative AI

Claude (Opus 5; Anthropic) was used during manuscript preparation for language editing and consistency checking, and to draft portions of the analysis code, which the authors reviewed and verified. It was not used to design the study, to collect or generate data, or to interpret results. The authors take full responsibility for the content.

## Data, Materials, and Software Availability

Data from Cohort 1, together with the analysis code, are available on the Open Science Framework at https://osf.io/8kah2. Cohort 2 is part of an ongoing longitudinal study and its data are not yet publicly deposited; the deidentified dyad-level measures supporting the analyses reported here are available from the corresponding author on reasonable request, subject to a data use agreement and Virginia Tech Institutional Review Board approval.

## Author contributions

Q.L. designed research; Y.-Y.C. performed research; J.K.-S., B.C., and Y.Q. contributed new reagents or analytic tools; Q.L. analyzed data; Q.L. and T.-H.L. interpreted the results and wrote the paper; and Y.-Y.C., J.K.-S., B.C., and Y.Q. edited the manuscript. All authors approved the final version.

## Competing interest statement

The authors declare no competing interest.

## Acknowledgments

We thank the former and current members of the Affective NeuroDynamics & Development Lab and the JK Lifespan Development Lab at Virginia Tech for their help with data collection. We are grateful to the adolescents and parents who participated in our study. This work was supported by a Virginia Tech Institute for Society, Culture and Environment research award to T.-H.L. and by the National Institute on Drug Abuse (Grant R01 DA036017) to J.K.-S.and B. C.

## Notes

### Competing Interest Statement

The authors have declared no competing interest.

