## Supplementary Materials for "Parent–Child Cortical Similarity Indexes Multiple Dimensions of Sensitivity in the Association Between Family Environment and Adolescent Internalizing Symptoms"

**SM1. Sample and exclusions**

Cohort 1. Seventy-six families were recruited. Fifty-one dyads had complete T1-weighted scans for both members and passed visual quality inspection; three dyads were subsequently excluded because of incomplete demographic or questionnaire data, yielding 48 dyads for the primary models. Cohort 2. Of 70 dyads drawn from an ongoing community-based longitudinal study, one lacked complete T1-weighted data, yielding 69 dyads with usable imaging that passed visual quality inspection of the FreeSurfer reconstruction; two of these were excluded for an incomplete attachment questionnaire, yielding 67 dyads for the primary models; dyads with missing data beyond this were excluded rather than imputed, because dyadic similarity cannot be meaningfully imputed from one member’s data. Some Cohort 1 families contributed more than one dyad. The 48 Cohort 1 dyads comprise 48 different children and 47 different parents from 43 families: 39 families contributed one dyad, three contributed two dyads each (a sibling pair, with each child paired to a different parent), and one contributed three dyads (three siblings, one paired with the mother and two with the father, which is why 48 dyads involve 47 rather than 48 parents). Nine of the 48 children (18.8%) had a sibling in the sample, and no child was paired with both parents. Cohort 2 comprised 67 dyads from 67 families, so the 115 analytic dyads come from 110 families in total. Standard errors are clustered by family in every model, and the cross-family permutation null excludes any pairing of a child with a parent from their own family, including a sibling’s parent; across the 51 Cohort 1 dyads with complete imaging this yields 2,538 rather than 51 × 50 = 2,550 cross-family pairs, whereas Cohort 2 contained no siblings (69 × 68 = 4,692 pairs). Sample characteristics are given in Table 1; the two cohorts differed in household income, parental education, proportion of mother–child dyads, and child age, and did not differ in racial composition (predominantly White in both). The cohorts also differed in recruitment frame: Cohort 1 families were recruited as a healthy community sample, whereas Cohort 2 was drawn from a community sample enriched for substance-use risk. Symptom severity in the clinical range was therefore assessed in Cohort 2 only (Table 1), and the two cohorts are not compared on it.

**SM2. Image acquisition and head-size control**

Cohort 1 was scanned on a 3T Siemens PRISMA scanner with a 64-channel matrix head coil at the Fralin Biomedical Research Institute at Virginia Tech Carilion (T1-weighted: TR = 2.5 s, TE = 2.06 ms, flip angle = 8°, 1 mm isotropic voxels, FOV = 256 mm). Cohort 2 was scanned on a 3T Siemens Tim Trio scanner with a 12-channel head coil (T1-weighted: TR = 1.2 s, TE = 2.66 ms, flip angle = 8°, 1 mm isotropic voxels, FOV = 245 × 245 mm, 192 slices). Participants were excluded if they did not meet MRI safety criteria, including claustrophobia, head injury with loss of consciousness exceeding 10 minutes, orthodontia impairing image acquisition, severe psychopathology, and other contraindications. All T1-weighted images were processed with FreeSurfer 8.1.0 using the standard recon-all pipeline, and cortical reconstructions were inspected visually before analysis.

Each parcel–feature value was residualized on estimated total intracranial volume (eTIV) before any similarity calculation. The regression was fitted leave-one-family-out: for each family, coefficients were estimated on all other families, and the family’s residuals were standardized against the training residuals. This prevents a family from contributing to its own reference distribution, which would induce similarity by construction.

**SM3. Family environment harmonization**

Cohort 1 used the Family Environment Scale (FES) (1) and Cohort 2 the Inventory of Parent and Peer Attachment (IPPA) (2). The primary pairing was FES Relationship with IPPA Alienation (reverse-scored), selected because both index the warmth-to-hostility quality of the parent–child bond; higher scores indicate a more positive family environment in both cohorts. Valence checks confirmed that all subscales correlate in the expected directions with the internalizing outcome in both cohorts; these correlations were recomputed on the analytic dyads (*n* = 48 and 67), matched by scan identifier.

**SM4. Outcome variables**

Internalizing symptoms were the first principal component of five z-standardized indicators, extracted separately within each cohort because the instruments differ. Cohort 1: perceived stress (PSS), depressive symptoms (CES-D), emotion dysregulation (DERS-18), negative affect (PANAS-NA), trait anxiety (STAI-T); PC1 = 75.4% of variance. Cohort 2: perceived stress (PSS), depressive symptoms, anxiety symptoms, negative affect (PANAS-NA), emotion dysregulation; PC1 = 50.8% of variance. In Cohort 1, participants missing three or more indicators of five were excluded; remaining missing values were multiply imputed (m = 20, chained equations via scikit-learn IterativeImputer, max_iter = 20, sample_posterior = True), with the final score the mean of the 20 imputed PC1 scores. In Cohort 2, all analytic participants had complete indicator data and PC1 was extracted from complete cases. Three loading checks (sign, version, and identifier match) are run whenever the outcome is loaded to guard against silent file errors.

**SM5. Standardization**

Similarity was computed after three standardization steps, all applied before any parent–child comparison and all serving one requirement: the resulting score must express similarity within a family rather than similarity any two brains share by construction. Throughout, let x(i, v, m) denote the value of morphometric feature m at parcel v for participant i, with v = 1, …, 400 (Schaefer parcels) and m = 1, …, 5.

*Step 1*: head-size removal (leave-one-family-out). For each family f, a linear regression of each parcel–feature value on estimated total intracranial volume (eTIV) was fitted using all participants from the other families: x(i, v, m) = α(v, m) + β(v, m)·eTIV(i) + ε. The coefficients estimated without family f were then applied to the members of family f, *r*(i, v, m) = x(i, v, m) − α̂₍₋f₎(v, m) − β̂₍₋f₎(v, m)·eTIV(i), and the residuals were scaled by the standard deviation of the training-set residuals. Fitting leave-one-family-out prevents a family from contributing to its own reference model, which would otherwise induce resemblance by construction (SM2).

*Step 2*: within-subject standardization. Each participant’s residualized values were z-scored across the 400 parcels, separately per feature: z(i, v, m) = [*r*(i, v, m) − mean_v *r*(i, ·, m)] / SD_v *r*(i, ·, m). After this step a participant is described by the shape of their cortical profile rather than its overall level, following the within-subject normalization used in morphometric similarity mapping (3).

*Step 3*: removal of the population-average pattern (leave-one-family-out). Because all human cortices share a common regional architecture, two unrelated brains are already highly similar; this shared pattern was removed by standardizing each parcel–feature value against its distribution across individuals from all other families, computed within cohort: d(i, v, m) = [z(i, v, m) − μ₍₋f₎(v, m)] / σ₍₋f₎(v, m). The resulting deviation scores express how each individual departs from the group-typical pattern, so that parent–child similarity quantifies whether two family members depart from that pattern in the same way. For phenotypic similarity this step is applied to parcel values; for architectural similarity it is applied to network edge weights after network construction (SM6). That these steps preserve a family-specific component is shown by the comparison of own-parent similarity with every other family’s parent (Fig. 2B; Table S5c).

**SM6. Parent–child cortical similarity measures**

Five morphometric features were extracted from each Schaefer-400 cortical parcel: cortical thickness, surface area, gray-matter volume, Gaussian curvature, and mean curvature. Parent–child cortical similarity was quantified in two forms — phenotypic and architectural — using the standardized morphometric values described in SM5.

*Phenotypic similarity.* Phenotypic similarity quantified parent–child correspondence in regional morphometric profiles. For a cortical unit (whole cortex, cortical system, or individual network), parent–child similarity was calculated separately for each of the five morphometric features as the Pearson correlation between the parent’s and child’s standardized parcel-wise values across all parcels:

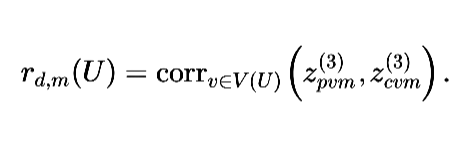

This yielded five feature-specific correlations for each dyad and cortical unit. The primary phenotypic score was a composite obtained by Fisher -transforming these correlations, averaging across the five features, and transforming the mean back to the correlation scale:

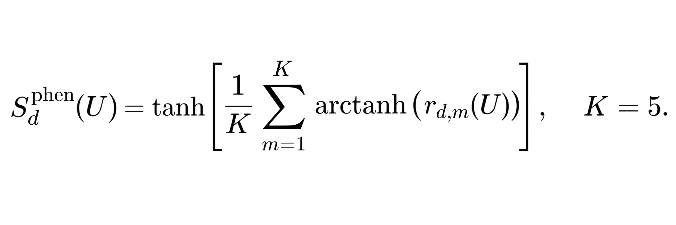

Thus, phenotypic similarity reflects correspondence between parent and child in the spatial distribution of regional morphometric characteristics. Feature-specific scores were retained for exploratory analyses, whereas the composite score was used in the primary whole-cortex and system-level analyses.

*Architectural similarity.* Architectural similarity quantified parent–child correspondence in interregional morphometric organization. Following Seidlitz et al. (3), a morphometric similarity network (MSN; see also 4) was first constructed separately for each participant from the within-participant standardized values produced in Step 2 of SM5. For each pair of parcels, the MSN edge was defined as the Pearson correlation across the five morphometric features:

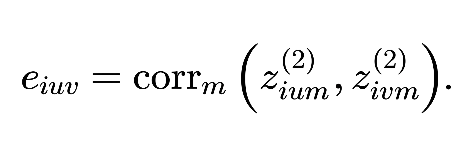

This procedure yielded a symmetric matrix for each participant, comprising 79,800 unique edges for the whole cortex. After construction of the MSNs, population-average edge patterns were removed using the leave-one-family-out standardization described in SM5.

For each cortical unit, architectural similarity was then calculated as the Pearson correlation between the parent’s and child’s standardized edge vectors, restricted to edges whose two endpoints:

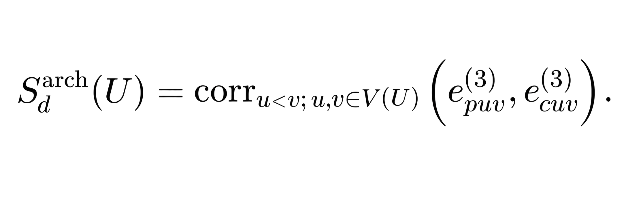

Architectural similarity therefore reflects correspondence in how cortical regions are organized relative to one another, rather than correspondence in the regional morphometric values themselves. Morphometric similarity networks have been linked to cytoarchitectonic organization and structural connectivity (3, 5).

*Relation between the two forms and cortical units.* Phenotypic and architectural similarity are derived from the same morphometric measurements but summarize different aspects of cortical organization. Phenotypic similarity directly compares corresponding regional values, whereas architectural similarity compares the pattern of interregional relations derived within each participant. Consequently, similar phenotypic similarity scores need not imply similar architectural similarity scores, and vice versa.

The same procedures were applied at the whole-cortex, system, and network levels. System-level phenotypic scores were computed directly across all parcels assigned to that system, and system-level architectural scores were computed directly across all within-system edges; system scores were not obtained by averaging network-specific similarity estimates.

**SM7. Cortical partitions**

The primary division groups the visual, somatomotor, and limbic networks as a sensory–limbic system and the remaining four networks as an association system. The limbic network’s placement departs from functional hierarchies, on which limbic cortex ranks toward the association end: when the archetypal sensorimotor–association axis (6) is parcellated with the Schaefer-400 atlas, the limbic network has a mean rank of 281 of 400, between the frontoparietal and default mode networks (network assignment via the 17-network Schaefer-400 labels mapped onto the seven-network solution, so parcel counts differ slightly from those used here). It is grouped with the sensory networks because paralimbic and neocortical cortex follow distinct developmental programs across adolescence, with morphometric similarity increasing in paralimbic cortex while decreasing in neocortical areas (7). Two alternative divisions defined in advance were tested: the visual and somatomotor networks alone, following the tethering account of Buckner and Krienen (8), and the limbic network alone. Results under every division are reported in Table S4b.

**SM8. Statistical framework**

Pooled models combined dyads after within-cohort z-standardization of all continuous variables, with interaction terms built from standardized variables, a cohort indicator in every model, and standard errors clustered by family. Covariates were the parent–child age difference, child race (White versus non-White contrast), and sex pairing (four levels, daughter–mother reference). Meta-analytic models were estimated per cohort and combined by fixed-effect inverse-variance weighting, with Cochran’s Q and *I*² reported descriptively. Differences between forms were tested as δ = β(architectural × E) − β(phenotypic × E) from a single model containing both scores, with the standard error from the full covariance matrix. The difference between systems in that contrast was tested in the model containing all four system-by-form scores as Δ = δ(sensory–limbic) − δ(association), using the contrast vector (−1, +1, +1, −1) over the four interaction coefficients ordered sensory–limbic phenotypic, sensory–limbic architectural, association phenotypic, and association architectural, with the standard error from the full covariance matrix of those four coefficients. Because the four scores are partialled against one another, the δ values in that model differ from those estimated in the two-form models (Table S1). Specificity contrasts compared the own-parent score with each control (placebo-parent similarity; child atypicality, the norm of the child’s standardized deviation profile) in the same manner. The pre-specified hypothesis family for specificity comprised the contrasts on scores carrying an interaction, corrected by Benjamini–Hochberg. The primary model adjusts for the parent–child age difference, child race (White versus non-White), and sex pairing, with a cohort indicator in every model. The child’s own age is not included: both it and the parent–child age difference index developmental variation, and the age difference is the term defined on the pair, which is the level at which the similarity scores are measured. Because the choice between them is not settled, two further models were estimated — one adding the child’s own age, and one adding the child’s own age with child race omitted. All three are reported in full in Table S2a and are treated as sensitivity analyses of the same data rather than as independent tests of the primary claims.

SM9. Regional heritability of parent–child correspondence

*Rationale.* Parent–child cortical similarity could in principle reflect shared genotype rather than shared family experience. Under additive transmission, the expected parent–child correlation at a region is proportional to that region’s heritability. This yields two things. The first is a check on the similarity measure: if the correspondence maps carry familial signal, they should align spatially with genetic architecture. Such alignment is expected under any account with a genetic component and does not separate genetic from shared-environmental contributions; it establishes only that the measure behaves as a measure carrying familial signal should. The second is a prediction from the genetic account itself: if the moderation reflected genetic transmission alone, it should be concentrated in the most heritable cortex. Family specificity is a separate question, tested by the unrelated-adult comparisons of Fig. 2 and Table S5c rather than here. The first analysis is reported in the main text; the second could not be evaluated at this sample size and is reported here for completeness.

*Heritability map.* Regional SNP-heritability for cortical thickness, surface area, volume, mean curvature and Gaussian curvature was taken from Table S29 of ref. 9 (UK Biobank and ABCD meta-analysis, *N* up to 36,843), which reports 180 bilaterally averaged regions of the HCP-MMP1 parcellation (10). Values were mapped to the 400 Schaefer parcels on fsaverage as the vertex-weighted average of the HCP-MMP1 regions each parcel overlaps; one parcel without coverage (7Networks_LH_DorsAttn_Post_17) was excluded, and homologous left and right parcels received the same value. As a check on the mapping, the projected values reproduce the enrichment patterns reported in ref. 9: thickness heritability is concentrated in idiotypic and somatomotor cortex, whereas surface-area heritability follows a distinct, paralimbic-weighted pattern. Ref. 9 leaves the regional phenotypes unadjusted for global measures, on the grounds that such adjustment biases regional estimates (Supplementary Note 3 of ref. 9), whereas the similarity scores used here have global effects removed (SM5). The mismatch is conservative for the present comparison: unless the global component itself tracks family-specific correspondence, it adds variance to one side of the correlation without adding covariance, and so attenuates rather than inflates the association.

*Positive control.* For each feature, the parent–child correlation across dyads was computed at every parcel from the standardized deviation values of SM5 (the 120 dyads with complete imaging, 51 and 69 by cohort) and correlated across parcels with the feature-matched heritability map. Significance used a parcel-centroid spin test (11) (5,000 rotations of the fsaverage sphere, right hemisphere mirrored); spin *P* values are the proportion of rotations at least as extreme as the observed correlation, and all spatial tests used the 399 parcels with heritability coverage. All five features were positive: surface area ρ = 0.30 (spin *P* = 0.001), gray-matter volume 0.15 (0.036), Gaussian curvature 0.10 (0.057), cortical thickness 0.11 (0.074), mean curvature 0.10 (0.099). The five-feature composite, which is the pre-specified quantity for all primary analyses (SM6), gave ρ = 0.25 (*P* = 0.001); single-feature maps carry only 12–32% true-signal variance, so the composite is the better-powered estimate and the feature-wise values are descriptive. For architectural similarity, the per-parcel average of the parent–child edge correspondence gave ρ = 0.21 (*P* < 0.001).

*Sensorimotor–association axis.* Heritability covaries with the sensorimotor–association axis (ρ = −0.18 for surface area to −0.54 for cortical thickness; five-feature mean −0.48), so the correspondence was recomputed with parcel-wise S-A rank partialled out. The association was undiminished: composite partial ρ = +0.28, surface area +0.32, architectural +0.22 (all spin *P* = 0.001); single features other than surface area fell to 0.09–0.13 (*P* = 0.05–0.11). The correspondence is therefore not a restatement of the S-A axis. Regional measurement reliability nonetheless bounds both quantities — less reliably measured parcels attenuate both the heritability estimate and the parent–child correlation — so part of the spatial alignment may reflect shared dependence on measurement precision rather than shared biology. The partial correlations control the dominant axis of this variation but cannot exclude it entirely, and a parcel-wise reliability map was not available in these cohorts.

*Moderation.* The phenotypic score decomposes exactly into parcel-wise contributions and the architectural score into edge-wise contributions (verified to numerical precision), so similarity can be recomputed over any subset of parcels or edges on the same scale. For the architectural score, each edge was assigned a heritability weight equal to the geometric mean of the five-feature mean heritability of its two endpoint parcels. This weight is a heuristic proxy rather than an edge-level heritability estimate: an edge is a correlation between two parcels across features, so its heritability depends on the joint genetic structure of the pair, including the genetic correlation between them, for which no published regional estimates exist. The node-level positive control above supports the proxy only in part, and cortical similarity networks are now known to carry common-variant genetic effects of their own, organized along cortex-wide gradients (12); edge-level genetic maps would therefore afford a sharper partition of the architectural score than the endpoint-based weights used here. Continuous weighting by heritability was the planned analysis but was abandoned because the two weighted scores correlated above 0.9 (VIF 10–20). Tertile splits within each system (VIF 1.3–1.7) gave, for sensory–limbic phenotypic similarity, β = 0.156 (*P* = 0.089) over the most heritable third and 0.204 (*P* = 0.030) over the least; for association architectural similarity, 0.163 (*P* = 0.095) and 0.256 (*P* = 0.002). The tertile split is not equally interpretable across the five features. For cortical thickness the heritability tertiles largely recode network identity within the sensory–limbic system: the most heritable third has a mean sensorimotor–association rank of 45 against 201 for the least heritable third, and the least heritable third contains all 26 limbic parcels. For surface area the two thirds are balanced on that axis (101 versus 101). The composite contrast therefore inherits part of this confounding; the composite is retained because it is the pre-specified quantity (SM6), not because the split is feature-invariant. The low-minus-high differences were δ = +0.07 (95% CI −0.28 to 0.42) and +0.14 (−0.12 to 0.40), neither larger than under 1,000 spin-rotated heritability maps (*P* = 0.64 and 0.46; 1,000 rather than 5,000 rotations because the moderation model is refitted at every rotation). These contrasts are uninformative by construction: with *SE*(δ) ≈ 0.15, detecting a difference with 80% power requires |δ| ≈ 0.42, roughly twice the entire interaction being partitioned (β = 0.22). Correspondingly, the coefficient over the least heritable third did not exceed the distribution obtained from spatially matched random thirds (0.172 ± 0.056 and 0.158 ± 0.076; *P* = 0.28 and 0.11), so no third of the cortex was distinguished. As a further check, each full-unit similarity score was residualized, at the dyad level, on the score computed from the most heritable third of parcels (edges, for the architectural level), and the standardized residual was entered as the moderator in the standard Brain × Environment model; the reported coefficient is the residual score’s interaction with family environment. The residualized coefficients were smaller than the unrestricted estimates but did not approach zero: sensory–limbic phenotypic β = 0.168 (*P* = 0.071) against β = 0.22 unrestricted, and association architectural 0.141 (0.196) against 0.22. The remaining unit-by-form cells were, for the phenotypic composite, whole cortex 0.046 (0.638) and association 0.086 (0.345), and for the architectural score, whole cortex 0.171 (0.060) and sensory–limbic 0.046 (0.574). Across the five features of the sensory–limbic phenotypic composite the residualized coefficients ranged from −0.003 to 0.240 and only gray-matter volume reached significance (0.240, *P* = 0.004; surface area 0.193, 0.063; cortical thickness 0.141, 0.253; mean curvature −0.023, 0.791; Gaussian curvature −0.003, 0.977). These estimates are not independent of the tertile contrasts above, carry the same power limitation, and are reported for completeness rather than interpreted. The analysis does not adjudicate the genetic alternative in either direction and is not interpreted in the main text.

**Supplementary tables**

Sample characteristics appear as Table 1 in the main text and are not duplicated here. Table S1. Main models: each unit-by-form similarity score and its interaction with family environment. Table S2a. The six primary cells under all three covariate specifications. Table S2b. Per-cohort estimates beside the pooled estimates. Table S2c. Correlations among the four system-by-form similarity scores and collinearity diagnostics. Table S3a. Dyadic specificity at the system level against unrelated-adult similarity and child atypicality. Table S3b. Dyadic specificity at the carrier networks (post hoc). Table S4a. Homogeneity of network-level interactions within each system. Table S4b. Alternative cortical divisions and dyadic specificity under each. Table S5a. Every network in both forms, and the difference between them. Table S5b. The full exploratory grid of seven networks by seven similarity measures. Table S5c. Per-network parent–child similarity against a cross-family permutation null. Table S5d. Whole-cortex family specificity: the values plotted in Fig. 2C and 2D.
